# GABAergic signaling by VIP interneurons gates running-dependent visual recovery in the adult brain

**DOI:** 10.1101/2025.05.21.655402

**Authors:** Anna Lebedeva, Friedrich Kling, Benjamin Rakela, Michael P. Stryker, Jennifer Y. Sun

## Abstract

Experience-dependent plasticity in the adult visual cortex is enhanced by locomotion, a process mediated by vasoactive intestinal peptide (VIP)-expressing interneurons. While VIP interneurons are known to signal through both Gamma-aminobutyric acid (GABA) and VIP peptide, the specific contributions of these pathways during different forms of plasticity remain unclear. Monocular deprivation (MD) in adult mice alters cortical responses, though more slowly and differently than during a critical period in early life. Here, we used two-photon calcium imaging in awake adult mice to dissect the roles of VIP and GABA release from VIP interneurons during adult MD and subsequent binocular recovery. We found comparable level of ocular dominance shifts after MD in mice deficient in either peptidergic or GABA signaling, but disrupting GABA signaling impaired recovery of binocular responses. We also showed that running preferentially enhances contralateral eye responses in binocular primary visual cortex. However, this eye-specific modulation of visual responses by running was altered during recovery from MD and was dependent on VIP signaling pathways. These findings highlight the GABA-mediated inhibition by VIP interneurons as a critical pathway for promoting visual restoration in the adult brain.

**Significance Statement:** Using longitudinal two-photon imaging in awake adult mice with genetically altered signaling pathways in VIP interneurons, we demonstrate that GABAergic, but not peptidergic, signaling from VIP interneurons is essential for the recovery of binocular vision following monocular deprivation. We further reveal that locomotion modulates cortical responses in an eye-specific manner, a property dynamically reshaped by plasticity and dependent on VIP interneuron function. These findings identify a discrete inhibitory circuit element that links behavioral state to sensory recovery and highlight GABA release from VIP cells as a potential therapeutic target for restoring visual function in adulthood.

## 1 Introduction

Experience-dependent plasticity, the remarkable ability of the brain circuit to undergo structural and functional changes in response to external stimuli, is a fundamental process for sensory and motor function maturation, learning, and memory formation [1, 2]. In the visual system, a robust model to study experience-dependent plasticity is ocular dominance plasticity (ODP) in the primary visual cortex (V1): during a tightly regulated time window, termed a critical period, brief monocular visual deprivation (MD) by eyelid suture causes a dramatic shift in ocular dominance, reducing responses to the closed eye and increasing responses to the open eye [3]. This change in neuronal responses is accompanied by drastic shrinkage of thalamocortical axonal projections representing the deprived eye [4, 5]. Such ODP is a well-preserved visual developmental process across mammalian species, varying only in time specific to their maturation speed [5–8].

Compared to the robust plasticity during the critical period, the mature visual cortical circuit in the adult brain exhibits a more limited capacity for ODP [9–11]. In the mature V1, prolonged MD in adults is required to shift ocular dominance, which largely results from potentiated responses to the non-deprived eye rather than decreased responses to the deprived eye [12–14]. Many studies have examined effective ways to enhance ODP in the adult visual cortex, including directly perturbing the excitation-inhibition balance with pharmacological intervention [15–17], early life MD experience [18] or a period of light deprivation [19] preceding MD, inhibitory neuron transplantation in the adult brain [20], and long-term environmental enrichment [21, 22]. Further studies demonstrate running, even just during the MD period can promote ODP in the adult mouse visual cortex [23, 24], and vasoactive intestinal peptide (VIP)-expressing interneurons have been identified as critical players in this process: optogenetic activation of VIP interneurons alone, rather than general aerobic exercise, is sufficient and necessary for enhancing cortical plasticity in adult mice [24].

VIP interneurons are a unique population of cortical cells that secrete both the inhibitory neurotransmitter gamma-aminobutyric acid (GABA) and the VIP, and provide preferential inhibition to other types of interneurons in the cortex [25–27]. VIP interneurons are activated during locomotion [28], which provide an overall disinhibitory effect onto pyramidal neurons and lead to increased visual responses during running [29]. This increased neural activity contributes to increased synaptic integration and multiple forms of cortical plasticity in the adult brain [30, 31]. Vasoactive intestinal polypeptide, a 28-amino-acid polypeptide initially isolated from gastrointestinal nerves, [32] is found highly concentrated in the VIP cells of rodent cerebral cortex [33–35]. The peptide has been suggested to be a modulator of synaptic function, increasing cyclic AMP formation through its synergistic interaction with norepinephrine [36, 37], thereby increasing spontaneous neural firing [38, 39]. VIP signaling has also been associated with various cognitive processes, including spatial learning, fear conditioning, and social behavior [40–43].

Previous studies have established the importance of VIP interneurons in mediating the enhancement of visual cortical plasticity induced by running in the adult brain [24], but the specific contributions of their GABAergic and peptidergic signaling pathways remain unclear. In this study, we aimed to further dissect the respective roles of the two pathways in facilitating ODP promoted by locomotion. To that end, we used two transgenic mouse lines to prevent either VIP or GABA release from VIP interneurons (Fig 1A). In the VIP-KO (referred as VIP^−/−^) animals, the production of vasoactive intestinal polypeptide in all VIP neurons was disrupted by deleting the peptide-encoding genes globally [44]; since VIP interneurons are the sole source of the peptide in the brain, the global knockout is also a specific deletion from VIP cells. In the VIP-cre;Vgat^*fl/fl*^ (referred as Vgat^−/−^) animals, Cre-mediated recombination specifically disrupts GABA secretion from VIP interneurons by deleting the vesicular GABA transporter [45]. We used two-photon calcium imaging to track the ocular dominance changes in the primary visual cortex of awake, head-fixed animals when they underwent MD and subsequent binocular recovery. We found that while both transgenic lines displayed ocular dominance plasticity, the recovery of binocular vision during running-promoted ODP was compromised in the mice in which GABAergic secretion from VIP interneurons was disrupted. The facts that the initial ocular dominance shift did not require the inhibition produced by VIP interneuron, but that the normal recovery from MD did so, highlights the critical role of VIP-cell mediated inhibition in the recovery of function promoted by locomotion in the adult brain.

**Figure 1:**
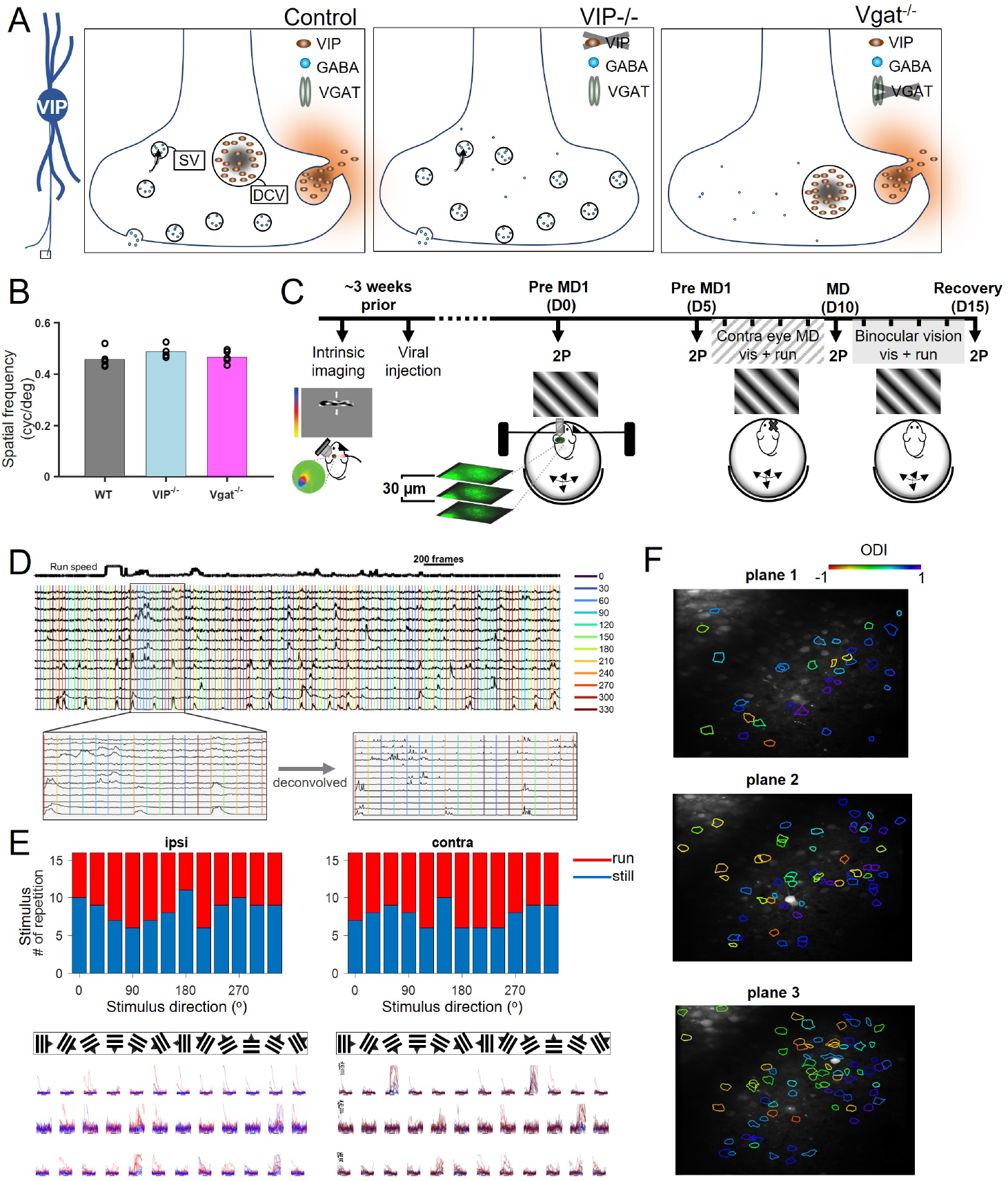
Chronic imaging of animals going through MD and recovery. **A)** Illustration of release of VIP peptide and GABA by VIP neurons. Two transgenic lines VIP^−/−^ and Vgat^−/−^, where the VIP or GABA signaling pathway was disrupted, respectively. **B)** VIP^−/−^ and Vgat^−/−^ animals exhibit normal visual acuity comparable with control (CT) animals. The highest spatial frequency gratings that elicited an optomotor reflex were calculated by taking the average responses between the two eyes. **C)** Experimental timeline: binocular V1 was identified using intrinsic signal imaging for viral injection of GCaMP6s. Two-photon calcium imaging was conducted at 4 time points when animal went through monocular deprivation and recovery. **D)** Example of calcium dynamics in a control animal. Drifting gratings of 12 orientations were randomized and shown to either eye using pneumatic eye shutters in awake, head-fixed animal. Running speed was monitored through the process and plotted at the top of the calcium traces. **E)** Three example cells are shown, with the responses to each orientation shown in red and blue for running and still trials. **F)** An example session with 3 planes images, and ROIs (neurons) are identified and labeled with colored contours denoting the ODI values.

## 2 Results

### 2.1 Measuring binocular visual responses in V1 with two-photon imaging

We first verified that the global disruption of VIP signaling pathways does not impair fundamental visual processing using a virtual optomotor system [46] to measure the optomotor reflex in both transgenic lines and their non-mutant littermate controls (referred as CT): the visual acuity was comparable among the three groups (0.458±0.032 cycles/° for CT, 0.488±0.0264 cycles/° for VIP^−/−^, and 0.467±0.0248 cycles/° for Vgat^−/−^ groups respectively, p = 0.28 ANOVA test, Fig 1B).

To assess experience-dependent changes in visual cortical responses, we applied two-photon calcium imaging to chronically track visually evoked activity in layers 2/3 neurons of the binocular zone in V1 in awake, head-fixed adult mice, while animals went through monocular deprivation and binocular recovery. The binocular zone of V1 was identified using intrinsic signal imaging [47]. The genetically encoded calcium indicator GCaMP6s [48] was expressed in binocular V1 either by injection of a CGaMP virus or in transgenic GCaMP animals [49]. After two baseline measurements of V1 responses to visual stimulation delivered independently to the two eyes, animals underwent 5 days of MD by suturing the lid of the eye contralateral to the recorded visual cortex, followed by 5 days of recovery after re-opening the lid of the deprived eye. Every day throughout the ten-day protocol of MD and recovery full-field sinusoidal drifting gratings were presented to the animals, which had been head-fixed in a custom apparatus that allowed them movement on a floating styrofoam ball (Fig 1C). Individual sessions of visual stimulation lasted 2-3 hours.

Mice were imaged under two-photon microscopy for visually evoked responses from the two eyes at four time points: pre-MD1 (Day 0), pre-MD2 (Day 5, just prior to lid suture), post-MD (Day 10), and after binocular recovery (Day 15). For each imaging session, two or three planes were taken of neurons at different depths within cortical layers 2/3 while the head-fixed mice were free to run or stand still [29]. Running speed was tracked continuously. Full-field sinusoidal gratings drifting in 12 directions were randomly interleaved and presented to the contralateral or ipsilateral eye using automated eye shutters. Each neuron’s ocular dominance index (ODI, which varies from +1 for neurons responding solely to the contralateral eye to −1 for pure ipsilateral eye responses) was determined by comparing the visually evoked responses to ipsilateral- and contralateral-eye stimulation at the neuron’s preferred orientation, and population responses were calculated from all neurons from all imaging planes (Fig 1D-F).

### 2.2 Defects of recovery from monocular deprivation in Vgat^−/−^ animals

To determine whether the running-promoted ocular dominance plasticity in adult animals remains intact in mice in which VIP peptide or GABA secretion is disrupted in VIP cells, we tracked the ODI in each group across four time points: day0 (pre-MD1), day 5 (pre-MD2) after which the contralateral eyelid was sutured, day 10 (5d MD) before which the eyelid was re-opened, and day 15 (recovery) (Fig 2A, also see the timeline in Fig 1C). The overall ODI at each time point was calculated by bootstrapping the same number of neurons from each animal within each group (see Method, Fig 2B). Consistent with previous findings, the control group exhibited a significantly decreased ODI after 5d MD (ODI = 0.28±0.04 for pre-MD1 and 0.30±0.04 for pre-MD2, 0.01±0.04 for 5d MD, estimated median with 95% confidence interval, N=5), indicating a shift in overall neuronal responses toward the ipsilateral eye, which had remained open. ODI also shifted significantly as a result of MD in the VIP^−/−^ and Vgat^−/−^ groups: VIP^−/−^:ODI =0.40±0.03 for pre-MD1 and 0.35±0.03 for pre-MD2, 0.22±0.04 for 5d MD, estimated median with 95% confidence interval, N=5; Vgat^−/−^:ODI = 0.39±0.03 for pre-MD1 and 0.34±0.03 for pre-MD2, 0.12±0.041 for 5d MD, estimated median with 95% confidence interval, N=4). No significant difference between the two baseline measurements were found in any of the three groups.

**Figure 2:**
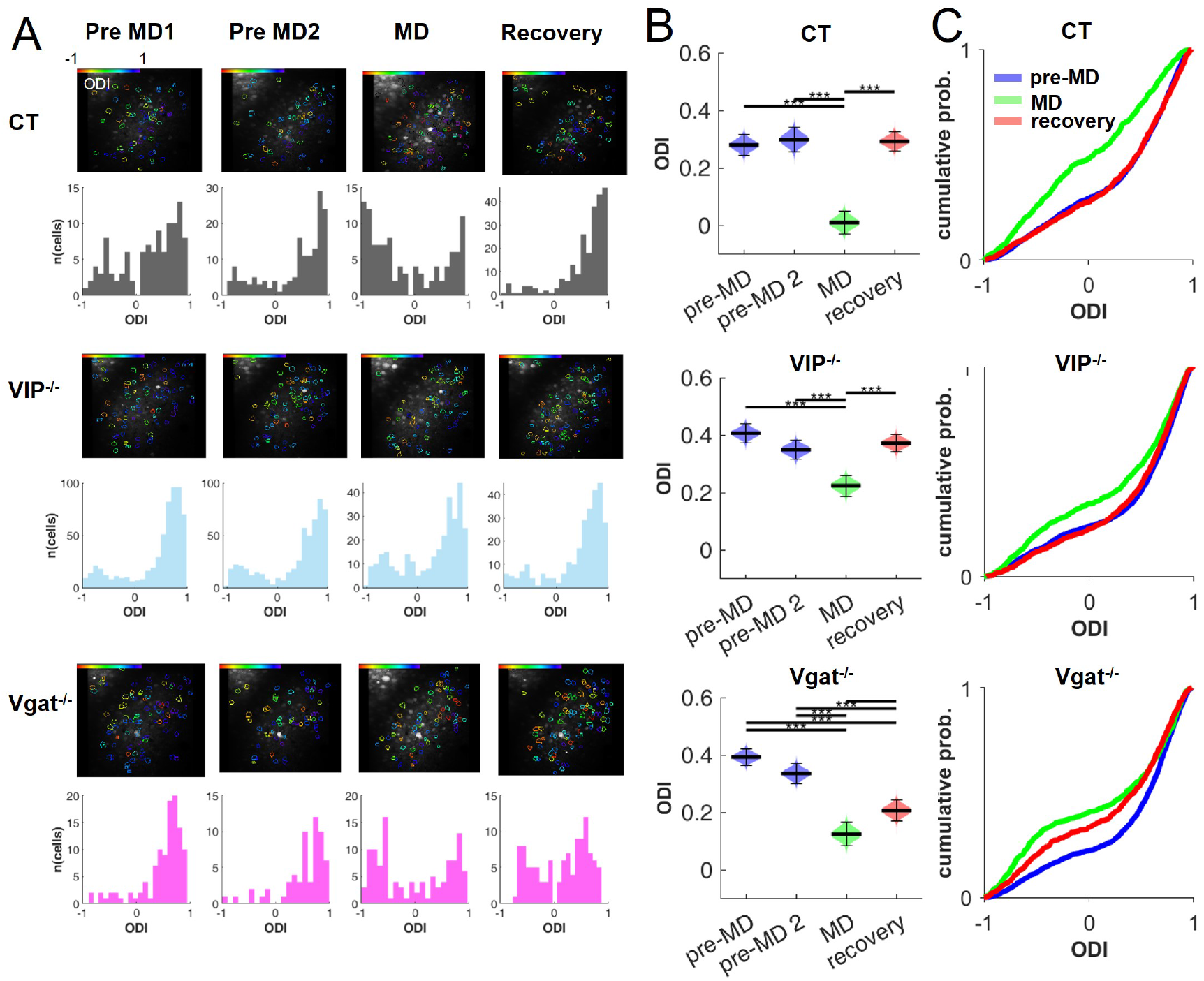
Ocular dominance plasticity and recovery was compromised in Vgat^−/−^ animals. **A)** Example longitudinal imaging of a matched field-of-view over 4 time points: pre-MD1(day0), pre-MD2 (day5), 5d MD (day10), and 5d recovery (day 15), for a control animal, a VIP^−/−^, and a Vgat^−/−^ animal. The 2-photon imaging is shown for only one plane, with all identified ROIs with outlined color representing their ODI (top panel). Neurons across all planes for each session are pooled to generate the histograms of ODI values (bottom panel). **B)** Summary of ODI values. Data are calculated by bootstrapping the same number of neurons from each imaging session on a given day, 10000 draws. P-value *p < 0.05, ** p < 0.005, *** p < 0.0005. No significant difference was detected between pre-MD and pre-MD 2 in any groups. **C)** CDF distributions of ODI values across stages, in comparison with baseline (blue line, pre-MD 1 and pre-MD2 combined as one pre-MD stage), 5d MD (green line) shifted the ODI towards more ipsilateral eye-responsive, but the 5d recovery (red line) in Vgat^−/−^ was incomplete.

After the 5 days of MD, animals experienced a recovery period of 5 days of binocular exposure with continued visual stimulation when head-fixed on a floating ball. The ODI in the control group rebounded to a level similar to that before MD (CT: ODI = 0.29±0.03, estimated median with 95% confidence interval). Responses to the formerly closed eye also recovered fully in the VIP^−/−^ animals (ODI = 0.37±0.03, estimated median with 95% confidence interval). In contrast, responses in the Vgat^−/−^ group recovered from MD only to a lesser extent (ODI = 0.21±0.03, estimated median with 95% confidence interval) that was significantly different from the baseline values prior to MD.

The failure of recovery in the Vgat^−/−^ animals is clearly illustrated in the cumulative distribution functions (CDF) of ODIs, by including all cells from all animals in each group, and combining the 2 baseline measurements to a single pre-MD distribution (Fig 2C). In both the control and the VIP^−/−^ groups, the ODI distributions pre-MD and following recovery are almost congruent (CT: p=1; VIP^−/−^: p=0.5, KS test with Bonferroni correction), and are widely separated from the distribution after MD (CT: p<10^−6^ for pre-MD vs MD, p<10^−6^ for MD vs recovery; VIP^−/−^: p<10^−6^ for pre-MD vs MD, p<10^−6^ for MD vs recovery; Kolmogorov-Smirnov test with Bonferroni correction). In the Vgat^−/−^ group, while MD shifted the ODI distribution towards the ipsilateral eye to a similar extent (p<10^−6^ for pre-MD vs MD, p = 1.0*10^−6^ for MD vs recovery, KS test with Bonferroni correction), the ODI distribution after the recovery period remained separated from the pre-MD distribution (p=0.001, KS test with Bonferroni correction). This result indicates that the functional recovery from MD promoted by visual stimulation during locomotion was compromised in animals in which GABA secretion from the VIP cells was disrupted.

### 2.3 Eye-specific running modulation is differentially impacted by binocular recovery in animals with disrupted VIP signaling pathways

In addition to its effects on long-term plasticity, running is also known to enhance neural activity and increase the gain of orientation-selective responses in mouse V1 [29], with the strongest modulatory effect in layers 2/3 neurons [50–53]. We asked whether the modulatory effect of locomotion on V1 responses is similar for stimulation delivered to the two eyes; whether the eye-specificity changes as a result of MD and recovery; and how it is affected by disruptions of the two signaling pathways of VIP cells.

To directly measure the effect of running on visually evoked responses, we focused on neurons that have similar numbers of running and still trials during visual stimulation presented to each of the two eyes (example cases in Fig 3A, see Methods). Consistent with previous reports, running enhanced visually evoked responses from both the ipsilateral and contralateral eyes, as indicated by the right-shifted CFD curves. The contralateral eye response was more significantly increased by running than the ipsilateral eye response.

**Figure 3:**
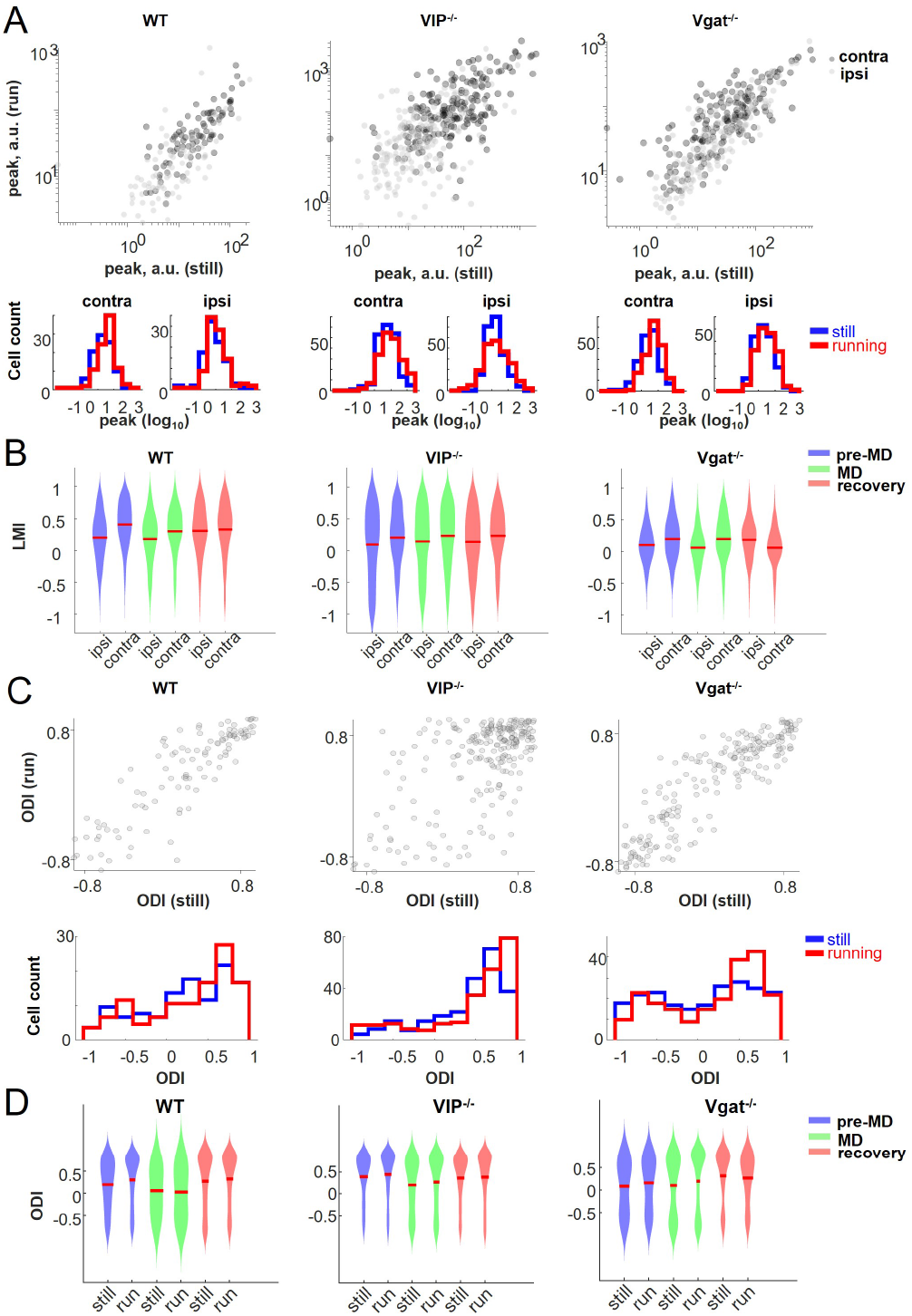
Running effect on neural responses are eye-specific. **A)** Running modulation of the visually evoked response for one example control animal (left), one VIP^−/−^ animal (center), and one Vgat^−/−^ animal (right), on baseline sessions. Top panels: averaged peak response depend on the behavioral state (running or still), with each point corresponding to one neuron. Middle panel: CDF distributions of the log values of the peak responses from the top plot, for each eye, during either running or still. Bottom panels: Histogram distributions of the peak responses from the top plot, separately plotted for each eye under various behavioral states. **B)** Summary of the locomotion modulation index (LMI) through different stages of ocular dominance plasticity, for the control animals (left), VIP^−/−^ animals (center), and Vgat^−/−^ animals (right). Data included all neurons that had some running and still trials, and the red line indicates mean. **C)** Example of ODI variation between running vs. still. Top panel: ODI in the same cell varies across the behavioural state, one example control animal (left), one Vip^−/−^ animal (center), and one ^−/−^ animal (right); one baseline session each. X-axis, still, Y-axis, running. Each point corresponds to one neuron. Bottom panel: distributions of the values of the ODI from the top plot, during either running or still. **D)** Summary of ODI comparisons between running and still conditions, for each animal group and each stage. Data included all neurons that had some running and still trials, and the red line indicates mean.

To systematically quantify the eye-specific running effect, we computed a locomotion modulation index (LMI) for each neuron, separately for their contralateral and ipsilateral responses (Fig 3B, see Methods). Indeed, running enhanced visually evoked responses for inputs from both eyes, but its modulatory effect on two eyes was consistently different, with a stronger effect on the contralateral eye responses at the baseline stage for all 3 animal groups (CT: ipsi, 0.20±0.01; contra, 0.40±0.01, mean±SEM, p=5*10^−25^ paired t-test, N = 800 cells from 7 sessions; VIP^−/−^: ipsi, 0.09±0.01; contra, 0.2±0.01, mean±SEM, p = 7*10^−6^ paired t-test, N = 794 cells from 6 sessions; Vgat^−/−^: ipsi, 0.09±0.01; contra, 0.19±0.01, mean±SEM, p = 7*10^−6^ paired t-test, N = 359 cells from 2 sessions). This difference between the effects of locomotion on the responses evoked through the two eyes remained after MD (CT: ipsi, 0.18±0.03, contra 0.29±0.03, mean±SEM, p=0.02 paired t-test, N = 114 cells from 3 sessions; VIP^−/−^: ipsi, 0.14±0.01; contra, 0.22±0.01, mean±SEM, 7*10^−4^ paired t-test, N = 757 cells from 5 sessions; Vgat^−/−^: ipsi, 0.05±0.01; contra, 0.19±0.01, mean±SEM, p = 10^−10^ paired t-test, N = 544 cells from 2 sessions).

The small number of imaging sessions in which animals had comparable numbers of running and still trials limits the power of comparisons of running-modulation across different time points. However, while the running modulatory effect for all three groups of animals was significantly stronger for the contralateral eye, and this eye-specificity persisted after monocular deprivation, it was no longer consistent after binocular recovery. While the VIP^−/−^ group retained the stronger LMI for contralateral eye (ipsi, 0.13±0.01; contra, 0.23±0.01, mean±SEM, p = 10^−5^ paired t-test, N = 728 cells from 5 sessions), in the control group, LMIs for for the two eyes became similar (ipsi,0.3±0.01; contra, 0.32±0.01 mean±SEM, p=0.39 paired t-test, N=483 cells from 4 sessions). In contrast, Vgat^−/−^ animals exhibited a higher LMI for ipsilateral eyes (ipsi, 0.18±0.02; contra, 0.05±0.02, mean±SEM, p = 0.0017 paired t-test, N = 135 cells from 3 sessions). These differences that emerged during recovery indicate that the circuit involved in running-modulation was significantly and differentially altered by ODP for animals with disrupted signaling pathways in their VIP cells.

### 2.4 Recovery from monocular deprivation is compromised in Vgat^−/−^ animals under both still and running states

The eye-specific modulation of responses by locomotion raises the possibility that ocular dominance might vary under different behavioral states. To address this, we calculated the ODIs separately for the same neurons in two different behavioral states: ODI_*run*_ using just running trials and ODI_*still*_ using only still trials (example cases in Fig 3C). Before MD, ODI_*run*_ was significantly higher than ODI_*still*_ in all three groups (Fig 3D purple line, CT: 0.31±0.01 vs 0.20±0.01, p =10^−17^ paired t-test, N = 800 cells from 7 sessions; VIP^−/−^: 0.44±0.01 vs 0.39±0.01, p =7*10^−4^ paired t-test, N=794 cells from 6 sessions; Vgat^−/−^: 0.16±0.01 vs 0.08±0.01, p = 3*10^−6^ paired t-test, N = 359 cells from 2 sessions). After 5d MD, the control group did not show a significant difference in ODI between running and still conditions (0.02 0.02 vs 0.05 0.02, p = 0.40 paired t-test, N=114 cells from 3 sessions). In contrast, VIP^−/−^ and Vgat^−/−^ groups continued to exhibit a significantly higher ODI_*run*_ than ODI_*still*_ (VIP^−/−^: 0.27±0.02 vs 0.20±0.02,p= 4*10^−5^, paired t-test, N=757 cells from 5 sessions; Vgat^−/−^: 0.20±0.02 vs 0.1±0.02, p = 2*10^−10^ paired t-test, N=544 cells from 2 sessions). This difference is consistent with LMI finding of Fig 3B, in which eye-specific modulation is less significant in the control group after 5d MD.

Interestingly, the difference between ODI_*run*_ and ODI_*still*_ became diverse between the three groups after binocular recovery: despite exhibiting similar eye-specific modulation, the control group returned to its original state after binocular recovery, with ODI_*run*_ higher than ODI_*still*_ (0.32 0.02 vs 0.27 0.02, p = 0.006 paired t-test, N=483 cells from 4 sessions). Following recovery from MD in the VIP^−/−^ group, ODI_*run*_ and ODI_*stil*_ became similar to each other despite the substantial eye-specific modulation by running (0.39±0.01 vs 0.37±0.01, p=0.16 paired t-test, N=728 cells from 5 sessions). In the Vgat^−/−^ group, ODI_*run*_ instead became significantly smaller than the ODI_*still*_ following recovery from MD (0.27±0.01 vs 0.32±0.02, p = 0.03 paired t-test, N=135 cells from 3 sessions). These results indicate that an animal’s behavioral states can affect the balance of responsiveness to the two eyes even when there is comparable modulation by running.

Therefore, we revisited the ocular dominance shift under the same behavioral states, first considering only trials during which the animals remained still (Fig 4A, B). Due to reduced data from the behavioral state criteria and our finding that no significant difference was detected between pre-MD and pre-MD 2 in any groups (Fig 2B), we combined the two pre-MD sessions to estimate baseline response. MD computed only from the still trials significantly shifted the ocular dominance towards the ipsilateral eye in all three groups (CT: 0.21± 0.06 for pre-MD, 0.09± 0.05 for MD, VIP^−/−^: 0.38± 0.05 for pre-MD, 0.14± 0.05 for MD; Vgat^−/−^: 0.36± 0.07 for pre-MD, 0.15 ± 0.09 for MD, estimated median with 95% confidence interval; p = 1.10^−6^ for CT, p < 10^−6^ for VIP^−/−^ and Vgat^−/−^, KS test with Bonferroni correction). Subsequent binocular recovery at least partially restored the preference towards the re-opened eye but to significantly different degrees in three groups (CT: 0.33±0.07, VIP^−/−^: 0.38±0.04; Vgat^−/−^: 0.21±0.07, estimated median with 95% confidence): While the control group had the strongest shift in ODI towards contralateral eye, its recovery was incomplete (p<10^−6^ for MD vs recovery, p<10^−6^ for pre-MD and recovery, KS test with Bonferroni correction). In contrast, the VIP^−/−^ group exhibited full rescue from the MD condition (p<10^−6^ for MD vs recovery, p = 0.10 for pre-MD and recovery, KS test with Bonferroni correction), yet the recovery in the Vgat^−/−^group did not recover to a level comparable to pre-MD (blue line) unlike the control or VIP^−/−^ group (p=4*10^−5^ for MD vs recovery, p = 0.0004 for pre-MD and recovery, KS test with Bonferroni correction).

**Figure 4:**
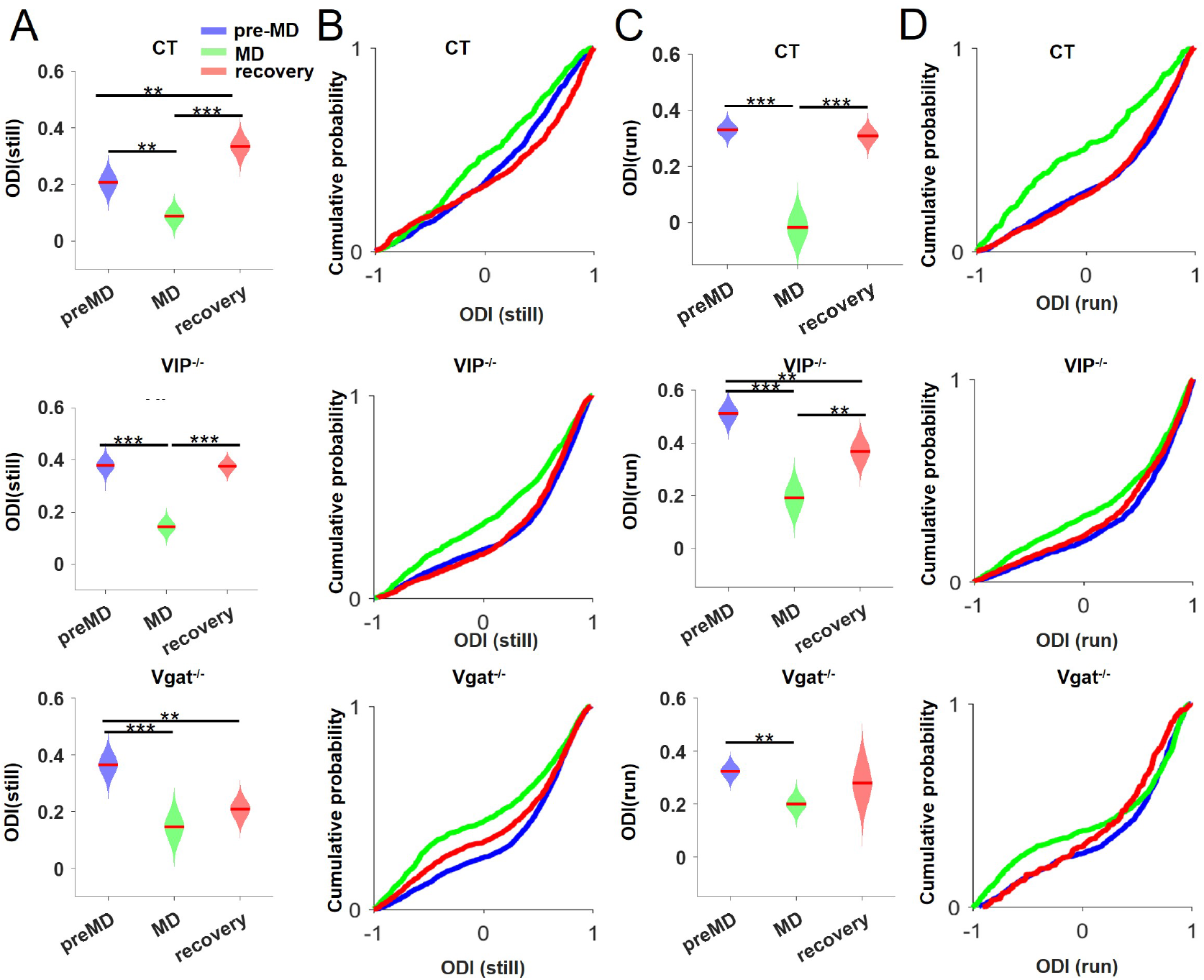
Recovery from monocular deprivation is compromised in Vgat^−/−^ group. **A)** Example of running modulation of the ODI. Top panels: modulation of ODI, depending on the behavioural state for one example control animal (left), a Vip^−/−^ animal (center), and a Vgat^−/−^ animal (right); one baseline session each. Each point corresponds to one neuron. Bottom panels: histogram distributions of the values of the ODI from the top plot, during either running or still. **B)** Summary of ODI comparisons between running and still conditions, for each animal group and each stage. **C)** CDF distributions of ODI values across MD conditions separately for still trials. Group CDFs were obtained by including all cells of each animal that has enough trials during still states. **D)** Similar as C, but in run trials. P-value *p < 0.05, ** p < 0.005, *** p < 0.0005

Analyzing only data from the running conditions (Fig 4C,D) produced similar findings. In comparison to control group, binocular recovery was less able to restore the ocular dominance towards preference for the re-opened contralateral eyes in VIP^−/−^ and Vgat^−/−^ groups (CT: p < 10^−6^ for MD vs recovery, p=0.42 for recovery vs pre-MD; VIP^−/−^: p =0.001 for MD vs recovery, p = 0.011 for recovery vs pre-MD; Vgat^−/−^: p= 0.008 for MD vs recovery, p=0.004 for recovery, KS test with Bonferroni correction). Together, these results revealed incomplete functional recovery from monocular deprivation in Vgat^−/−^ animals under both still and running states, supporting the idea that this process requires the intact GABAergic signaling in the VIP cells.

## 3 Discussion

As previously established, 5d monocular deprivation combined with daily visual exposure and running in the adult animals was sufficient to cause a shift of ocular dominance towards the open eye [24, 30]. In the present work, we applied 2-photon imaging to track neurons in layers 2/3 over the course of ODP and binocular recovery, in control animals and those with disrupted secretion of VIP peptide or GABA by theVIP cells. While VIP peptide plays a critical role in mediating many physiological functions in the brain circuits, mice lacking the VIP peptide (VIP^−/−^ mice) exhibited normal ocular dominance plasticity and binocular recovery, similar to the control group. In contrast, disrupted GABA release from VIP interneurons in Vgat^−/−^ mice led to a striking impairment in the recovery of binocular vision after monocular deprivation, whether or not animals were running during the recovery period. Overall, our findings highlight the critical involvement of VIP interneurons, particularly through GABAergic signaling, in mediating running-promoted recovery from MD in the adult visual cortex.

### 3.1 Eye-specific running modulation is altered during binocular recovery and requires GABAergic and peptidergic signaling in VIP interneurons

Locomotion is known to cause a global increase of V1 neural activity [28, 29, 50, 53, 54], our study further expands this finding by revealing that this modulatory effect is not uniform across inputs from the two eyes. Specifically, we show that running preferentially enhances contralateral eye responses in binocular V1, and that this eye-specific modulation is dynamically altered by experience-dependent plasticity. In control animals, the initial contralateral bias in running modulation disappeared after recovery, with responses to both eyes becoming similarly enhanced during running. This normalization suggests that the restoration of binocular vision involves a rebalancing of circuit dynamics, possibly through running-mediated neuromodulation. However, in mice lacking either GABA or VIP peptide release from VIP interneurons, this normalization fails to occur—or even reverses direction (e.g., increased ipsilateral bias in Vgat^−/−^ mice). These divergent outcomes imply that GABAergic and peptidergic signaling in VIP interneurons play essential and distinct roles in shaping state-dependent modulation.

While awake recordings are now routinely used to measure binocular responses [55–58], the current study, to our best knowledge, is among the first to report that the modulatory effect of running on V1 responses differs between the two eyes (see also [59]). This is not surprising given the running modulation on neural activity is not strictly multiplicative [53], and the input strengths from two eyes are different. Indeed, this points out the necessity to evaluate ocular dominance with consideration of effect of behavioral state.

### 3.2 Neither signaling pathways in VIP interneuron prevents running-promoted ocular dominance shift in the adult brain

Our findings demonstrate that neither the loss of VIP peptide (VIP^−/−^) nor the disruption of GABA release (Vgat^−/−^) from VIP interneurons prevents the initial ocular dominance shift induced in adult mice by MD combined with running. This suggests that running can promote competitive plasticity in the adult visual cortex through alternative neural circuits, even in the absence of intact VIP signaling pathways.

Previous studies have identified multiple interventions that enhance adult ODP, including environmental enrichment [60, 61], dark exposure [62], pharmacological reduction of intracortical inhibition [16, 63], and cross-modal sensory deprivation [64]. These manipulations likely alter the excitatory-inhibitory (E-I) balance in V1, creating a permissive state for plasticity. Our results align with this framework, as running—even in VIP^−/−^and Vgat^−/−^ mice — engages neuromodulatory systems, such as acetylcholine and norepinephrine, that may have more widespread effects and have been shown to facilitate cortical plasticity independently of VIP interneurons [2, 65, 66].

Alternatively, developmental compensation in VIP^−/−^ and Vgat^−/−^ mice may account for the preservation of the capacity for OD shifts. Since these mutant mice lack functional VIP peptide or GABA signaling from birth, other circuits may have adapted to maintain plasticity. Future studies using temporally precise manipulations could clarify whether acute disruption of VIP signaling affects MD-induced plasticity. For example, recent work by [67] demonstrated that acute blockade of peptidergic transmission using targeted conditional knockout creER system impairs specific forms of learning behavior. Such approaches may constitute novel, reliable tools for investigating neuropeptidergic systems in awake, behaving animals.

### 3.3 GABA release from VIP interneuron is essential for running-promoted recovery of cortical response following monocular deprivation

While both mutant groups exhibited normal ocular dominance shifts during the deprivation phase, only Vgat^−/−^ showed impaired recovery of responses to the deprived eye after binocular vision was restored. This selective deficit highlights a critical dissociation between the mechanisms driving initial deprivation-induced plasticity and those mediating functional recovery, with GABAergic signaling from VIP interneurons playing an indispensable role in facilitating running-promoted binocular vision recovery in the adult brain.

The failure of recovery in *Vgat*^−/−^ mice is consistent with prior work showing that eliminating VIP interneurons prevents the running-promoted visual enhancement in the adult mouse visual cortex [24]. Our current study further pinpoints the specific signaling pathway: while both GABAergic and VIP peptide are involved in plasticity, GABA release from VIP interneurons is the key mediator for the running-promoted plasticity. This is in line with recent work [68] showing that the locomotion-dependent stimulus-specific response enhancement of orientation-selective responses in adult visual cortex depends on GABA but not VIP release from VIP interneurons.

One plausible mechanism is that VIP-mediated disinhibition of pyramidal neurons—via suppression of SST and PV interneurons—alters the excitation-inhibition (E-I) balance during running, thus creating a permissive state for synaptic potentiation to enable stimulus-evoked strengthening of weakened synapses in the deprived eye. In the absence of GABA release from VIP interneurons, this disinhibitory “gate” remains closed even when the animal is running, preventing upregulation of deprived-eye inputs. This aligns with evidence that pharmacological blockage of NMDAR-dependent pathways prevents running-promoted enhancement in the adult brain [69].

### 3.4 Implications for human studies

Running appears to confer multiple benefits to the adult brain and to facilitate plasticity at multiple levels, by promoting neurogenesis, secretion of neurotrophic factors, and facilitating synaptic functions [70–73]. However, it remains controversial whether aerobic exercise directly increases V1 activity [74–76] or improves visual plasticity and learning in human subjects [77–79]. These discrepancies may arise from differences in exercise protocols, measurement techniques, or individual variability in neuromodulatory systems.

To date, the role of VIP interneurons in human visual plasticity remains unexplored. It is known that dysregulation of GABAergic cells is associated with pathophysiology underlying neurodevelopmental disorders[80], and VIP-positive interneurons are present in human visual cortex [81]. In mice, the dysfunction of VIP interneurons has been implicated in neurodevelopmental disorders characterized by altered sensory processing [82]. Our current finding corroborates the hypothesis that the plasticity-promoting effects of running in the adult brain are tightly linked to VIP interneuron-mediated disinhibition.

While the precise mechanisms remain to be elucidated, our findings point to VIP neuron-mediated GABAergic inhibition as a gating mechanism that links state-dependent cortical dynamics to recovery of sensory function. This mechanism may be particularly relevant for therapeutic strategies seeking to leverage behavioral interventions, such as exercise or visual training, to promote functional restoration in conditions like amblyopia or stroke.

## Methods

### Experimental Animals

All animal work was approved by the University of California San Francisco (UCSF) Institutional Animal Care and Use committee and conforms to the National Institutes of Health guidelines. VIP-cre;Vgat^*fl/fl*^ mice were generated by crossing the Vip-IRES-cre and Vgat^*fl/fl*^ mouse lines (stock number 010908 and 012897, the Jackson Laboratory). VIP-KO mice were acquired from Wascheck lab at UCLA (backcrossed 12 generations to C57BL/6J mice, Colwell et al., 2003). Littermates of each mutation line with normal functional VIP neurons were used as controls.

To allow calcium imaging, the calcium indicator GCaMP6s was expressed in all animals using one of two methods: some VIP-cre;Vgat^*fl/fl*^ animals and their control littermates were crossed with the Camk2a-tTA;tetO-GCaMP6s mouse line (stock number 024854 and 024742, The Jackson Laboratory) to label all excitatory neurons; the rest of mice (all VIP-KO mice and control littermates, as well as some VIP-cre;VGat^*fl/fl*^ animals) were injected with virus AAV2/1.hSynap.GCaMP6s.WPRE .SV40 (addgene #100843). Data for individual mice is listed in the table below. Mice were housed in the standard condition (12/12 dark–light cycle, free access to food and water) and adult mice (P90-150), both sexes of animals were used in the following study.

**Table 1:** Summary of experimental animals used in the study.

| **Group** | **ID** | **Genotype** | **Viral injection** |
| --- | --- | --- | --- |
| **Vgat<sup>-/-</sup>** | 6929 | VIP-cre(+/-);Vgat<sup>fl/fl</sup> | GCaMP6s |
| **Vgat<sup>-/-</sup>** | 6951 | VIP-cre(+/-);Vgat<sup>fl/fl</sup> | GCaMP6s |
| **Vgat<sup>-/-</sup>** | 6887 | VIP-cre(+/-);Vgat<sup>fl/fl</sup>;Camk2a-tTA;tetO-GCaMP6s |  |
| **Vgat<sup>-/-</sup>** | 6766 | VIP-cre(+/-);Vgat<sup>fl/fl</sup>;Camk2a-tTA;tetO-GCaMP6s |  |
| **VIP<sup>-/-</sup>** | 3360 | VIP-KO | GCaMP6s |
| **VIP<sup>-/-</sup>** | 3459 | VIP-KO | GCaMP6s |
| **VIP<sup>-/-</sup>** | 3477 | VIP-KO | GCaMP6s |
| **VIP<sup>-/-</sup>** | 3482 | VIP-KO | GCaMP6s |
| **VIP<sup>-/-</sup>** | 3487 | VIP-KO | GCaMP6s |
| **control** | 6871 | VIP-cre(+/-);Camk2a-tTA;tetO-GCaMP6s |  |
| **control** | 6884 | Vgat<sup>fl/fl</sup>;Camk2a-tTA;tetO-GCaMP6s |  |
| **control** | 3177 | VIP<sup>-/-</sup> littermate | GCaMP6s |
| **control** | 3223 | VIP<sup>-/-</sup> littermate | GCaMP6s |
| **control** | 3489 | VIP<sup>-/-</sup> littermate | GCaMP6s |

### Visual Acuity Test

Visual acuity of mice was measured using the virtual optomotor system (CerebralMechanics, Alberta, Canada; Prusky et al., 2004), consisting of four 17 inch LCD screens, simulating a rotating drum. An elevated platform was placed in the center of the virtual-reality chamber, and mice were placed onto it and allowed to adapt to the setup for 10 min on the first day before testing began. The mouse’s behavior was monitored using a video camera positioned above the platform. Vertical sinusoidal gratings of 100% contrast were presented on the screens at various spatial frequencies (0.042–0.514 cycles/degree) and randomized clockwise and counterclockwise directions, with a rotation speed of 12°/s. Head location was constantly tracked with manual click to ensure the correct presentation of the gratings. Successful detection is determined by smooth head movement in parallel with the rotation of the gratings, by experienced experimenters blind to the grating directions or mouse identities. At the end of each session, the highest spatial frequency that elicited noticeable tracking behavior was recorded as the threshold. Thresholds for the left and right eyes were determined separately with clockwise and counterclockwise rotation of the gratings, respectively. An average of those two values was taken as the representative threshold for each animal.

### Surgery and Intrinsic Imaging

One day before intrinsic imaging, a customized stainless steel plate for head fixation was attached to the skull with dental acrylic (Lang Dental Black Ortho-Jet powder and liquid) under isoflurane anesthesia (3% induction; 1.2-1.5% surgery). To evoke robust visually evoked responses, mice were injected with chlorprothixene (2 mg/kg, i.m.) and anesthesia was maintained using a low concentration of isoflurane (0.6 to 0.8% in oxygen); the core body temperature was maintained at 37.5°C using a feedback heating system. To identify binocular V1, the visual stimulus subtended 20° horizontal and was presented to one eye at a time (the non-tested eye blocked with an eye shutter) with the monitor positioned 25 cm directly in front of the animal. A temporally periodic moving 2° wide bar was generated using the Psychophysics Toolbox (Brainard, 1997; Kleiner et al., 2007) in Matlab (Mathworks) and continuously presented at a speed of 10°/s.

For chronic calcium imaging, a craniotomy was made over the identified binocular V1 in the left hemisphere (roughly 3 mm lateral to midline, 1 mm anterior to lambda) when animals were anesthetized with isoflurane (3% induction; 1.2-1.5% surgery), as well as a dose of subcutaneous injections of atropine (0.15 mg/kg), dexamethasone (1.5 mg/kg), and carprofen (15 mg/kg). In some animals, pAAV.Syn.GCaMP6s.WPRE.SV40 (addgene #100843-AAV1) was injected into three sites (50 nl/injection at 50 nl/min) in the binocular V1 at 200 and 350 m below the pial surface, using glass pipettes and a microinjection system (UMP3 UltraMicroPump, WPI). Finally, a 3-mm-diameter circular glass coverslip was secured using cyanoacrylate to allow long-term visualization of *in vivo* neuronal calcium activity.

### Monocular Deprivation and Binocular Recovery

Monocular deprivation was induced by suturing the right eyelid with a 7-0 polypropylene monofilament (Ethicon) under anesthesia. Suture integrity was inspected daily and immediately prior to each visual exposure session. Animals whose eyelids did not seal fully shut or had accidentally reopened were excluded from further experiments. After 5 days of deprivation, the suture was carefully removed under anesthesia.

### Visual Exposure

Visual stimuli were displayed on an LCD monitor (Dell, 30 × 40 cm, 60 Hz refresh rate, 32 cd/m^2^ mean luminance) placed 25 cm from the mouse (−20° to +40° elevation) with gamma correction. During two-photon imaging, drifting sinusoidal gratings of 3s at 12 evenly spaced directions (0.05 cycles per degree, and 1 Hz temporal frequency) were generated and presented in random sequence using the MATLAB Psychophysics Toolbox (Brainard, 1997; Kleiner et al., 2007), followed by 3s interstimuli interval of grey screen. During 5 days of MD and 5 days of recovery, animals were presented daily with randomized drifting gratings of 12 orientations for 2-3 hours while head-fixed to stand or run on the Styrofoam ball (Fu et al., 2015), with an average running time of 40.9 ± 13.6% (mean ± SD) during MD, and 48.1 ± 8.4% during binocular recovery.

### Two-Photon Calcium Imaging

After head-plate implantation mice were habituated to run or stand on a spherical treadmill (modified from the design of Dombeck et al., 2010, see also Fu et al., 2014) for high-resolution *in vivo* two-photon imaging. Running was monitored using optical mice; movement signals from the optical mice were acquired in an event-driven mode at up to 300 Hz, and integrated at 100 ms intervals and then converted to the net physical displacement of the top surface of the ball. A mouse was classified to be running on a single trial if its average speed for the 3s visual stimulus exceeds the running threshold (1 cm/s).

The imaging was performed using a resonant-galvo scanning, two-photon microscope (Neurolabware, Los Angeles, CA) and acquisition was controlled by MATLAB-based Scanbox software (Neurolabware). The light source is a mode-locked Ti:sapphire laser (Coherent Chameleon Ultra II) with excitation wavelength of 920 nm, green signals were detected through a 16× 0.8 NA microscope objective at 1.4-2.8 magnification. Images were acquired with a Nikon 16X water immersion objective (NA = 0.8, 3 mm working distance), in the layer 2/3 of the binocular visual cortex, located 150-310 m below the cortical surface (210 ± 50 m, mean ± SD) at a total sampling rate of 15.5 Hz (divided over 2 or 3 imaging planes, 30 m apart). Pupil size and ball movement were tracked using infrared light and high-speed cameras Dalsa Genie (teledyne Dalsa) with 740 nm long-pass filter during imaging. Imaging was acquired using neurovascular markers and customized software to maintain the same field of view over days.

To ensure comparable signals were exposed to both eyes, we used randomized visual exposures to either eye during the imaging session. During imaging, a pair of pneumatic eye shutters were positioned in front of each eye and randomly occluded one or the other eye before the visual presentation. Each eye of the animal was presented with visual stimuli of 12–16 repetitions of randomized drifting gratings (3 s duration) followed by 3 s interstimulus interval of blank screen (uniform 50% gray). Head-fixed mice were free to run on a spherical treadmill (air-supported polystyrene foam ball, diameter 20 cm) while viewing visual stimuli.

## Data Analysis

### Calcium Responses

Time-series imaging stacks were processed using Suite2p for motion correction, ROI (Region-of-interest) identification with customized classifier based on cell size and ellipticity, neuropil subtraction, and signal extraction, followed by manual verification. The slow calcium traces of each ROI was deconvolved using OASIS. Peak response for each trial was defined as the mean response during the 3 s of the visual stimulus, subtracted by the mean response during 1 s before the stimulus onset.

### Ocular Dominance Index

To calculate the ocular dominance index (ODI), the preferred orientation of each neuron was defined by the maximum peak response of either eye, regardless of either running or still conditions. Then, we calculated the mean response of the contralateral eye to the preferred orientation and to the opposite orientation (*response*_*contra*_). Then, we calculated the mean response of the ipsilateral eye to the preferred orientation and to the opposite orientation (*response*_*ipsi*_). The ODI was then calculated as the ratio between the difference of the contralateral and ipsilateral response, divided by their sum:

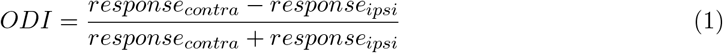

Similarly, we calculated ODI for running using only responses during running epochs and ODI for still using only the responses during still epochs.

### Session Selection

Running speed was calculated for the duration of visual stimulus representation (3 s), with the averaged speed greater than 1.5 cm/s considered running. For analysis requiring both running/still balance (Figure 3B,D), sessions with fewer than one running and one still trial for each of the 12 orientations and for each of the contralateral and ipsilateral eyes were excluded.

### Locomotion Modulation Index of Peak Response

Neural signals for each visual stimuli were first separated for running and still trials and averaged across trials. We calculated the peak response of the contralateral eye to the preferred orientation, during running and during still conditions. We then calculated the locomotion modulation index (LMI) for each eye:

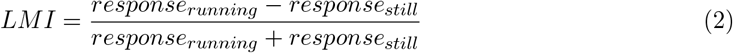

### Statistical Analysis

Statistical analyses were performed in Matlab (MathWorks). Statistical thresholds are indicated as \**p* < 0.05, \*\**p* < 0.01, \*\*\**p* < 0.001, ns: not significant.

To calculate the mean ODI for each time point, a non-parametric bootstrapping procedure was employed to equalize the impact of each animal on the estimation. The same number of cells were randomly sampled with replacement from each animal, with the number determined by the animal with the fewest passing cells. The bootstrap distribution of the mean ODI was generated for 10,000 iterations. From these distributions the mean was calculated.

To compare the ODI distributions, we recorded the differences of means between all combinations of groups upon each bootstrap iteration. We then performed a t-test of that distribution of difference against 0 to assess the significance of difference between ODI distributions.

Average cumulative density function (CDF) plots of mouse groups were generated as follows. First, for each group and day CDF distributions were calculated for individual animals. For each animal all ODIs of cells passing quality control were taken and CDF distributions generated with 2000 bins between −1 and 1. Group CDFs were obtained by averaging CDF distributions of animals in a group. Significant differences between CDF distributions were assessed using a bootstrapped KS test procedure: for each stage, the same number of cells were randomly sampled with replacement from each animal, with the number determined by the animal with the fewest passing cells. This was repeated 10,000 times to generate robust bootstrap distributions, followed by a KS test upon each draw to compare their empirical cumulative distribution functions between time points. For each test, the outcome was recorded positive if *p* < 0.05. The proportion of positive tests was then taken as the p-value.

## Acknowledgement

We thank Professor James Waschek of the University of California Los Angeles for the gift of the VIP-KO animals used in this study. This work was supported by NIH Grants K99-EY033976 (to B.R.), K99-EY029002 (to Y.J.S), and R01-EY02874 (to M.P.S.). Y.J.S and M.P.S. conceived the experiments, Y.J.S, A.L, and B.R. performed experiments, Y.J.S., A.L, and F.K. analyzed data, Y.J.S., A.L and F.K. wrote the manuscript, with editing by all authors.

## Notes

### Competing Interest Statement

The authors have declared no competing interest.

